# Epitope profiling reveals binding signatures of SARS-CoV-2 immune response and cross-reactivity with endemic HCoVs

**DOI:** 10.1101/2020.10.29.360800

**Authors:** Caitlin I. Stoddard, Jared Galloway, Helen Y. Chu, Mackenzie M. Shipley, Hannah L. Itell, Caitlin R. Wolf, Jennifer K. Logue, Ariana Magedson, Kevin Sung, Meghan Garrett, Katharine H.D. Crawford, Uri Laserson, Frederick A. Matsen, Julie Overbaugh

## Abstract

A major goal of current SARS-CoV-2 vaccine efforts is to elicit antibody responses that confer protection. Mapping the epitope targets of the SARS-CoV-2 antibody response is critical for innovative vaccine design, diagnostics, and development of therapeutics. Here, we developed a phage display library to map antibody binding sites at high resolution within the complete viral proteomes of all human-infecting coronaviruses in patients with mild or moderate/severe COVID-19. The dominant immune responses to SARS-CoV-2 were targeted to regions spanning the Spike protein, Nucleocapsid, and ORF1ab. Some epitopes were identified in the majority of samples while others were rare, and we found variation in the number of epitopes targeted by different individuals. We also identified a set of cross-reactive sequences that were bound by antibodies in SARS-CoV-2 unexposed individuals. Finally, we uncovered a subset of enriched epitopes from commonly circulating human coronaviruses with significant homology to highly reactive SARS-CoV-2 sequences.

## Introduction

A novel betacoronavirus, SARS-CoV-2, was transmitted into humans in late 2019 and has led to widespread infection, morbidity, and mortality across the globe (Wu et al., 2020). The disease caused by SARS-CoV-2 infection, Coronavirus disease 2019 (COVID-19), is characterized by a striking diversity in clinical presentation, ranging from asymptomatic or mild disease, to severe pneumonia and death. A number of studies have begun to address the role of the adaptive immune response in patients infected with SARS-CoV-2, but the repertoire of epitope targets linked to infection is only beginning to be comprehensively defined.

Coronaviruses (CoVs) have large, ~30 kb non-segmented genomes consisting of virus-specific accessory proteins and several universal open reading frames (ORFs): Spike (S), Membrane (M), Nucleocapsid (N), Envelope (E) and ORF1ab, which codes for a multitude of non-structural proteins (Chan et al., 2020; Cui et al., 2019). The S glycoprotein is recognized as highly immunogenic in SARS-CoV-2 infections, as well as for infections with the six other human CoVs (HCoVs) that are endemic and associated with the common cold (HCoV-OC43, HCoV-HKU1, HCoV-NL63, and HCoV-229E), or are the cause of more confined but highly pathogenic outbreaks in humans (SARS-CoV, MERS-CoV). The S protein of SARS-CoV-2 shares varying degrees of homology with other circulating CoVs ranging from 28% amino acid identity with the endemic HCoV-OC43 up to 76% with the highly virulent SARS-CoV (Walls et al., 2020; Wang et al., 2020). The S protein decorates the surface of all CoVs and mediates viral entry (Shang et al., 2020). Because of its surface exposure and role in infectivity, the S protein has been a major focus of vaccine development and recent efforts to isolate potent neutralizing antibodies targeting SARS-CoV-2 (Chi et al., 2020; Pinto et al., 2020).

While many vaccines are thought to protect by virus neutralization, antibodies that target viruses through mechanisms other than neutralization—often referred to as non-neutralizing antibodies—have been correlated with improved clinical outcomes for a variety of viruses, including HIV, influenza and Ebola (Lee and Kent, 2018; Mayr et al., 2017; Padilla-Quirarte et al., 2019; Saphire et al., 2018). Antibody responses to non-S CoV proteins have been detected previously, including non-neutralizing responses to the N protein of SARS-CoV, which is involved in genome packaging and is found in the mature virion core (Dutta et al., 2020). Interestingly, immune responses to the N protein of SARS-CoV-2 have recently been linked to poor clinical outcomes (Atyeo et al., 2020). Despite mounting evidence that the SARS-CoV-2 N protein may be highly antigenic in the context of COVID-19, there has been limited effort to fully characterize antibody responses mediated by N or the other non-Spike ORFs that are expressed during SARS-CoV-2 infection.

Sequence homology between SARS-CoV-2 and other circulating HCoVs increases the likelihood for cross-reactive antibody responses resulting from prior infection or vaccination. The N protein and other non-structural SARS-CoV-2 proteins are often more highly conserved than the S protein and thus may be targets for such cross-reactive non-neutralizing responses. Importantly, cross-reactive T-cell responses stemming from exposure to low-pathogenic endemic HCoVs have been identified in SARS-CoV-2 unexposed individuals (Grifoni et al., 2020; Mateus et al., 2020). Additional studies aimed at the B-cell immune response have identified cross-reactive antibody binding to the S proteins of SARS-CoV-2 and SARS-CoV, which share nearly 80% sequence identity genome-wide (Lv et al., 2020; Shrock et al., 2020). Despite lower degrees of homology than to the highly pathogenic SARS-CoV, cross-reactivity against the S protein from the four commonly circulating HCoVs in COVID-19 patient sera has also been identified (Shrock et al., 2020; Wölfel et al., 2020). Importantly, cross-reactive viral immune responses can be either cross-protective, as in the case of Influenza A and other viruses, or disease-enhancing, as in the case of dengue virus, and possibly SARS-CoV-2 (Arvin et al., 2020; Lee et al., 2020; Welsh et al., 2010). These divergent phenomena necessitate studies to evaluate SARS-CoV-2 antibody binding in unexposed individuals, and to measure sequence homology among high-binding epitopes from the full genomes of all HCoVs.

In order to capture the complete repertoire of neutralizing and non-neutralizing linear epitopes targeted by antibodies generated in the presence and absence of SARS-CoV-2 infection, we used a phage display immunoprecipitation approach (Larman et al., 2011) to profile immune responses in a population of patients with COVID-19 and patients with no exposure to SARS-CoV-2. We developed a pan-coronavirus (pan-CoV) phage library encompassing the complete proteomes of all human-targeted CoVs and used it to immunoprecipitate antibodies from samples from patients with mild or moderate/severe COVID-19, as well as SARS-CoV-2 unexposed patients, all collected in Seattle, Washington. Notably, we detected a pool of significantly enriched peptides from the S, N, and ORF1ab polypeptides of SARS-CoV-2 in samples from patients with COVID-19. We also identified four cross-reactive SARS-CoV-2 peptides that were enriched in pre-pandemic, SARS-CoV-2 unexposed individuals. Finally, we found a subset of endemic CoV peptides with high sequence conservation to enriched SARS-CoV-2 peptides in COVID-19 patients.

## Results

### A pan-CoV bacteriophage library detects antibody responses across the SARS-CoV-2 proteome

We generated a phage display library comprised of all seven CoVs known to infect humans, a bat SARS-like CoV, and a set of control peptides derived from the HIV-1 envelope sequence (**Fig. 1**). Oligonucleotide sequences were designed to cover complete CoV genomes in 39 amino acid tiles with 19 amino acid overlaps that were then cloned into T7 phage, amplified, and used in subsequent assays. This process was repeated to generate a replicate phage library from an independent oligonucleotide pool. We deeply sequenced both independent phage libraries and found similarly high coverage across the sequences included in the libraries, with > 98% of expected sequences detected (**Fig. S1A**).

**Figure 1.**
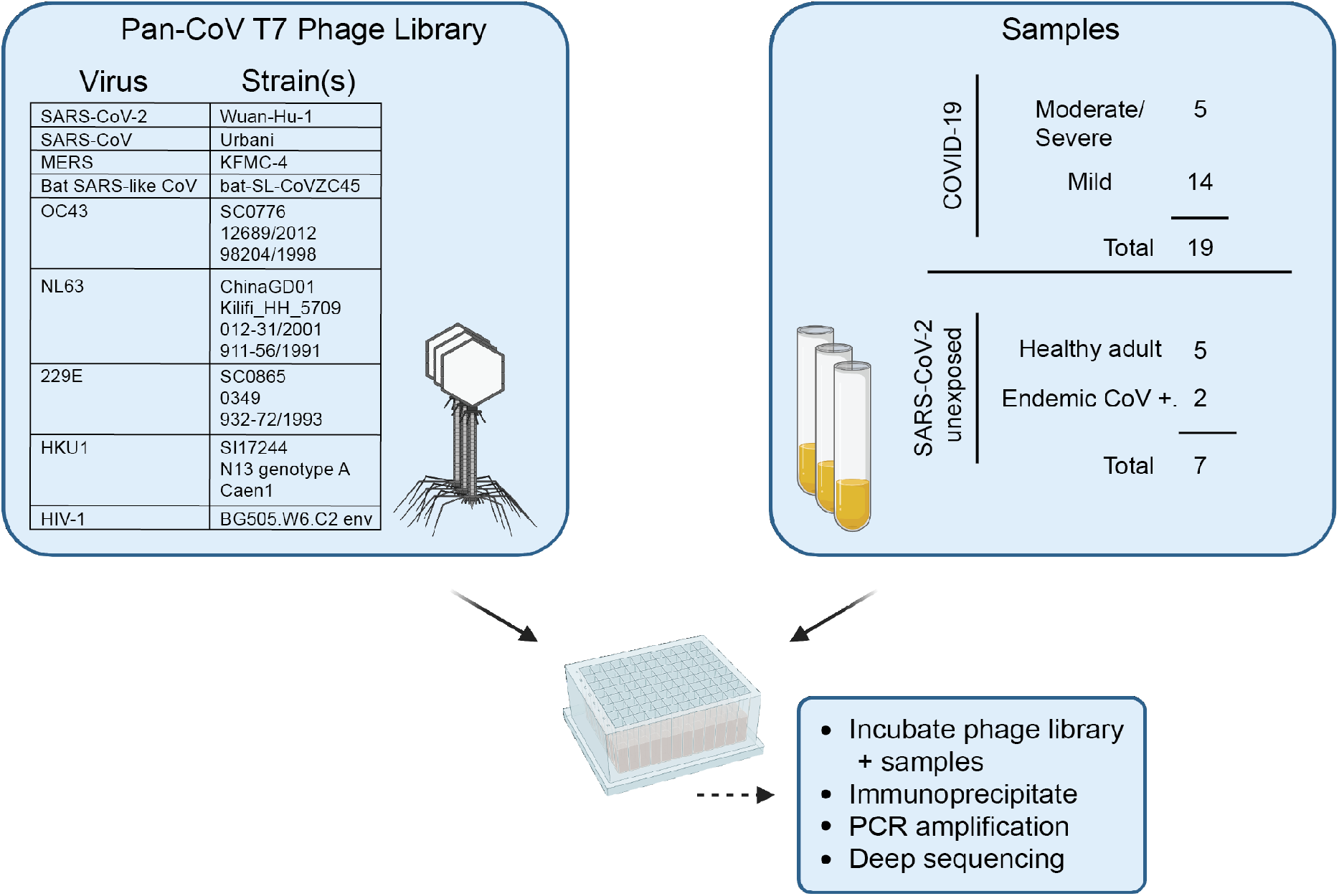
Summary of development of the pan-CoV T7 phage library and sample screening. Left panel, virus species and strains that comprise the pan-CoV phage library used in the study are listed. Right panel, summary of samples from COVID-19 or SARS-Cov-2 unexposed patients. The pan-CoV phage library and samples were combined in a plate-based immunoprecipitation assay and phage DNA was isolated for downstream sequencing and analysis. (p.s.o., post-symptom onset). Additional sample information can be found in **Tables 1 and S1**.

A total of 19 plasma or serum samples from patients with either mild or moderate/severe laboratory-confirmed SARS-CoV-2 infections (termed COVID-19 patients) was collected in Seattle, Washington as part of the Hospitalized and Ambulatory Adults with Respiratory Viral Infections (HAARVI) study or as residual clinical samples from hospital labs in Seattle. Samples from patients with mild COVID-19 cases were collected at approximately 30 days post symptom onset, and all moderate/severe samples were collected between 8-33 days post symptom onset. Samples from patients with endemic (non-SARS-CoV-2) HCoV infections were collected as part of the HAARVI study or were residual clinical samples from Harborview Medical Center (Seattle, Washington, USA). Archived samples collected from Seattle individuals before the pandemic were used as unexposed SARS-CoV-2 negative samples. Prior to study initiation, the following institutional human subjects review committee approved the protocol: University of Washington IRB (Seattle, Washington, USA), and concurrent approvals were obtained from the Fred Hutchinson Cancer Research Center. Additional sample demographic information is found in **Tables 1 and S1**.

**Table 1.**
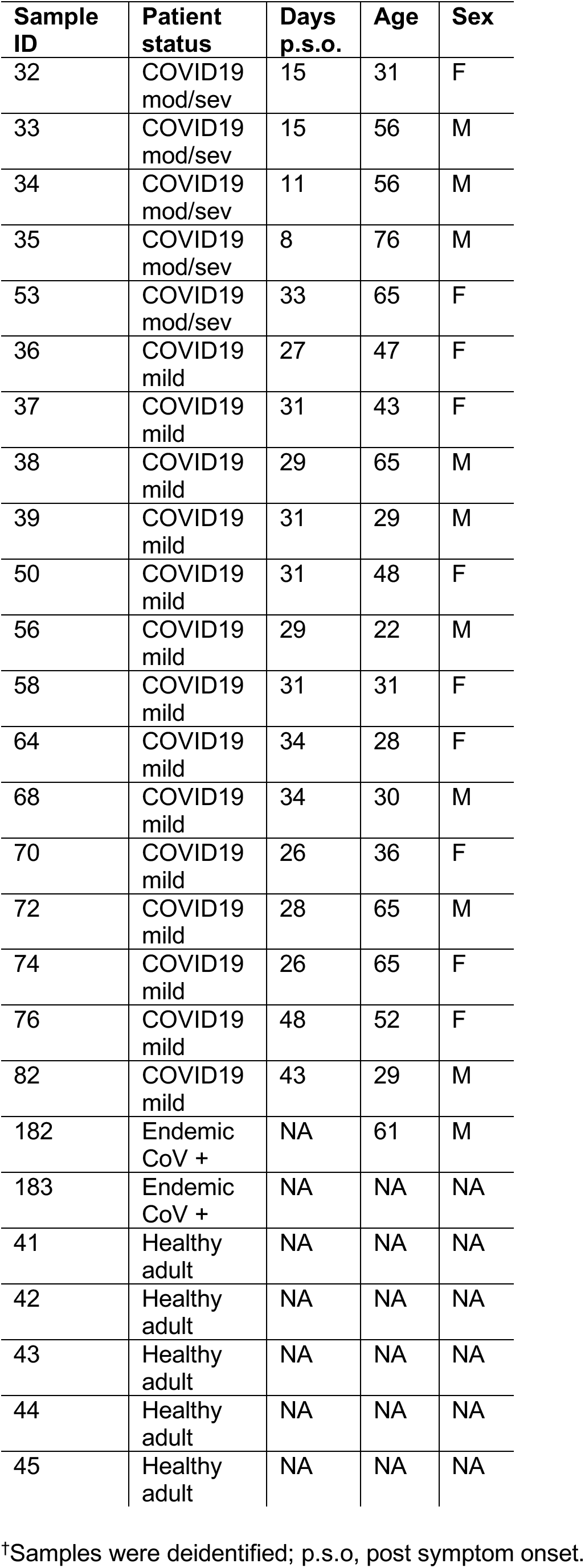
Sample information ^†^.

All samples were tested in a phage-based immunoprecipitation assay in which DNA from phage/antibody complexes was PCR amplified, and multiplexed samples were deep sequenced to determine enrichment of individual CoV peptides. We applied the following criteria to determine which samples to include in our downstream analyses: (1) a sample must have had a pairwise cross-correlation of at least 0.5 (Pearson’s R) between two technical (within-assay) replicates, and (2) a sample must have satisfied condition (1) in experiments conducted with both of the independent batches of the phage library (Libraries 1 and 2, **Fig. S1A**).

We performed a qualitative assessment of the SARS-CoV-2 epitope profile in COVID-19 patients by examining counts per million (CPM) from all SARS-CoV-2 ORFs. We detected signal for epitopes derived from all possible ORFs, but found significantly higher magnitude in S, N, and ORF1ab (**Fig. 2**). Signal was also detected for ORF3a and M, but at a much lower magnitude, with even lower signal for peptides enriched from the other proteins (note scale differences in **Fig. 2**). In order to evaluate the significance of epitope enrichment quantitatively, we modeled enrichment for all peptides from all samples along with a pool of mock-immunoprecipitation samples to account for non-specific peptide binding. We fit peptides to a Gamma-Poisson model, in which each sample-peptide pairing was assigned a minus log10 p-value (mlxp) (**Fig. 3A**). We exploited the HIV-1 envelope sequences in the pan-CoV libraries to estimate the false positive rate (FPR) of non-specific binding peptides. Next, we determined the mlxp cutoff corresponding to a FPR of 0.05 and identified 2,689 and 4,604 sample-peptide pairs, or “hits”, from phage Libraries 1 and 2, respectively (**Figs. 3A and S1B**). Across the two replicate experiments, there were 933 intersecting “hits”, 456 of which were unique peptides, where non-unique peptides were a result of sequence overlap between CoVs.

**Figure 2.**
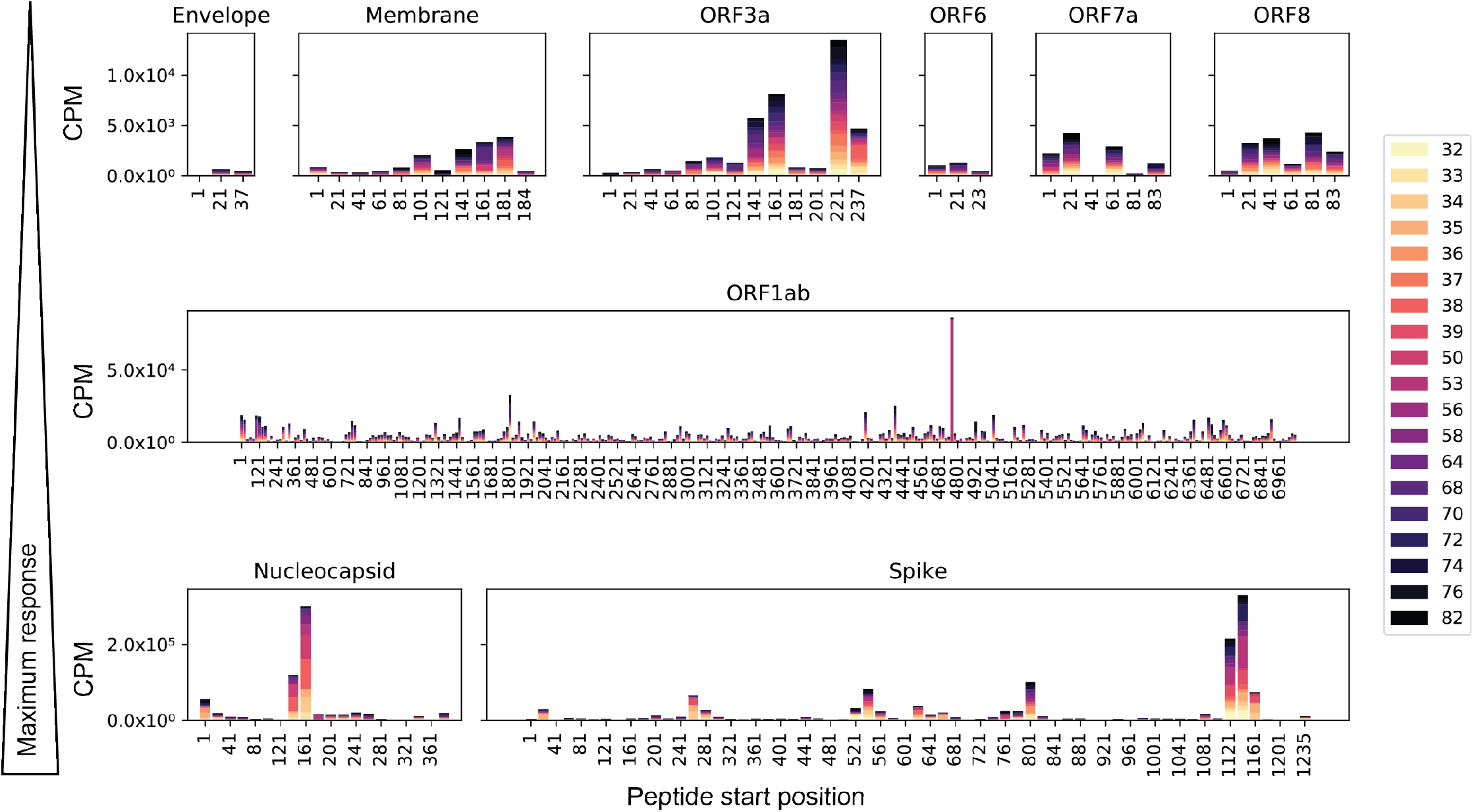
SARS-CoV-2 peptide enrichment based on raw counts per million. Individual panels showing enrichment for specific peptides among all COVID-19 patient samples along the lengths of nine SARS-CoV-2 ORFs. Panel rows are in order of increasing maximum response from top to bottom. Note the scales also increase in each row, indicating higher enrichment of the identified peptides. Bars are segmented by color for each sample included in the analysis, as depicted in the legend.

**Figure 3.**
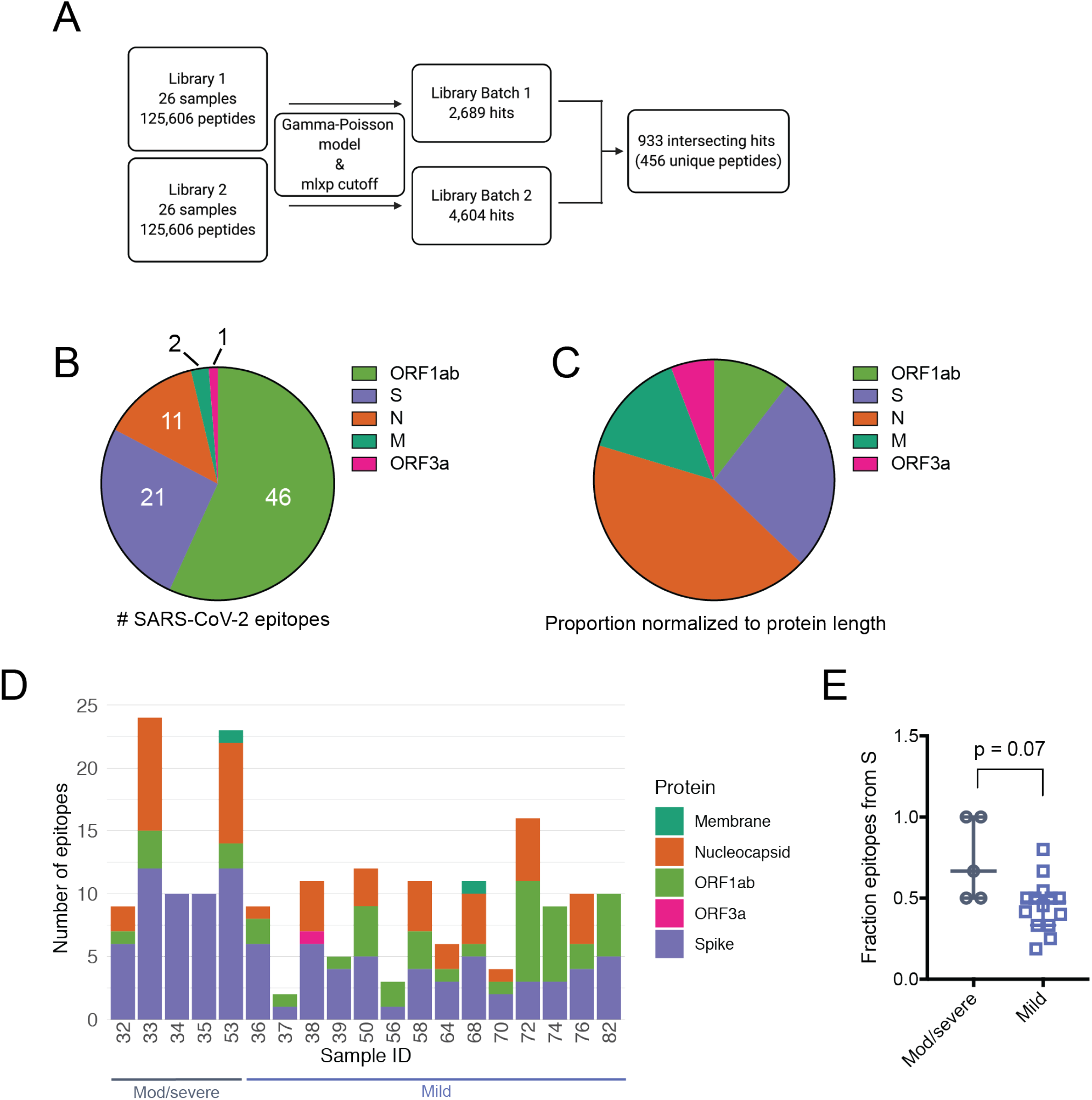
Results from global fit of all sample-peptide pairs with applied mlxp cutoff. (**A**) Data processing scheme. Samples were tested with two separate phage libraries (Library 1 and Library 2, **Fig. S1**). Peptide enrichment was scored using a Gamma-Poisson model and data were curated using a cutoff corresponding to FPR of 0.05 (**Fig. S1**). (**B**) Proportion of SARS-CoV-2 epitopes derived from individual proteins in all patient samples tested. Numbers indicate the total enriched SARS-CoV-2 epitopes from each ORF. (**C**) Proportions in (B) normalized with respect to polypeptide length. (**D**) Epitope counts across COVID-19 patient samples for SARS-CoV-2 only. Bars are further sectioned by SARS-CoV-2 ORF, indicated to the right. (**E**) Fraction of total epitopes arising from the S protein, calculated for moderate/severe and mild samples (# S epitopes/ # total epitopes). P value was calculated using a two-tailed unpaired Welch’s t test.

### Patterns of immune response at epitope resolution in COVID-19 patients and SARS-CoV-2 unexposed individuals

After fitting enriched immunoprecipitated epitopes, we detected significant responses in just five of the nine SARS-CoV-2 ORFs (**Fig. 3B**). We identified the most reactivity in ORF1ab, but after normalizing the number of reactive sequences by the ORF length, we found that the S and N proteins were dominant (**Fig. 3C**). Sparse responses to SARS-CoV-2 peptides were also detected within the M protein, and ORF3a (**Figs. 3B and 3C**). COVID-19 patient samples displayed variability in abundance of reactive epitopes, ranging from 2-24 reactive peptides in a given sample (**Fig. 3D**). All COVID-19 patient samples were reactive to epitopes from the S protein, while two moderate/severe COVID-19 samples, both less than twelve days post symptom onset, were reactive solely to the S protein and no other proteins (**Fig. 3D**). The proportion of the total epitope response arising from the S protein was higher in the five moderate/severe samples than in the mild samples (P = 0.07, Welch’s unpaired t test), suggesting that antibodies targeting the S protein are dominant in cases of moderate/severe COVID-19 (**Fig. 3E**).

We next examined the epitope profiles of all samples (both COVID-19 and SARS-CoV-2 unexposed) within the three immunodominant antigens: S, N, and ORF1ab. Within the S protein, we identified three key regions in which significant signal was detected, and these three regions were detected across the majority of individuals (**Fig. 4A and Table 2)**. The most dominant epitope was S_1121-1179 (composed of two adjacent peptides in our library that overlap by 19 amino acids), which spans a portion of the second heptad repeat (HR2) in the S2 subunit. In total, 84% (16/19) of COVID-19 samples were reactive to this epitope. The second region, S_801-839, spans the fusion peptide (FP) and includes the S2’ cleavage site. This region was reactive in 78% (15/19) of the COVID-19 samples. The third region, S_541-579 is located at the C-terminal end of the S1 subunit, immediately after the receptor binding domain (RBD) and upstream of the S1/S2 cleavage site. This epitope was reactive in 68% (13/19) of SARS-CoV-2 positive samples. Interestingly, S_541-579 was reactive in 100% (5/5) of the moderate/severe COVID-19 samples tested, but only 64% (9/14) of mild samples (**Figure 4A**). While not statistically significant within our current sample size (P = 0.68, Fisher’s exact test), this result may be suggestive of a correlation between COVID-19 severity and epitope patterning that should be explored in larger studies examining correlates of disease. Finally, we identified 13 additional dispersed S protein epitopes in smaller subsets of the individuals tested, including a sequence spanning the S1/S2 cleavage site (S_661-699), suggesting antibody recognition of pre-processed S protein (**Table 2**).

**Table 2.**
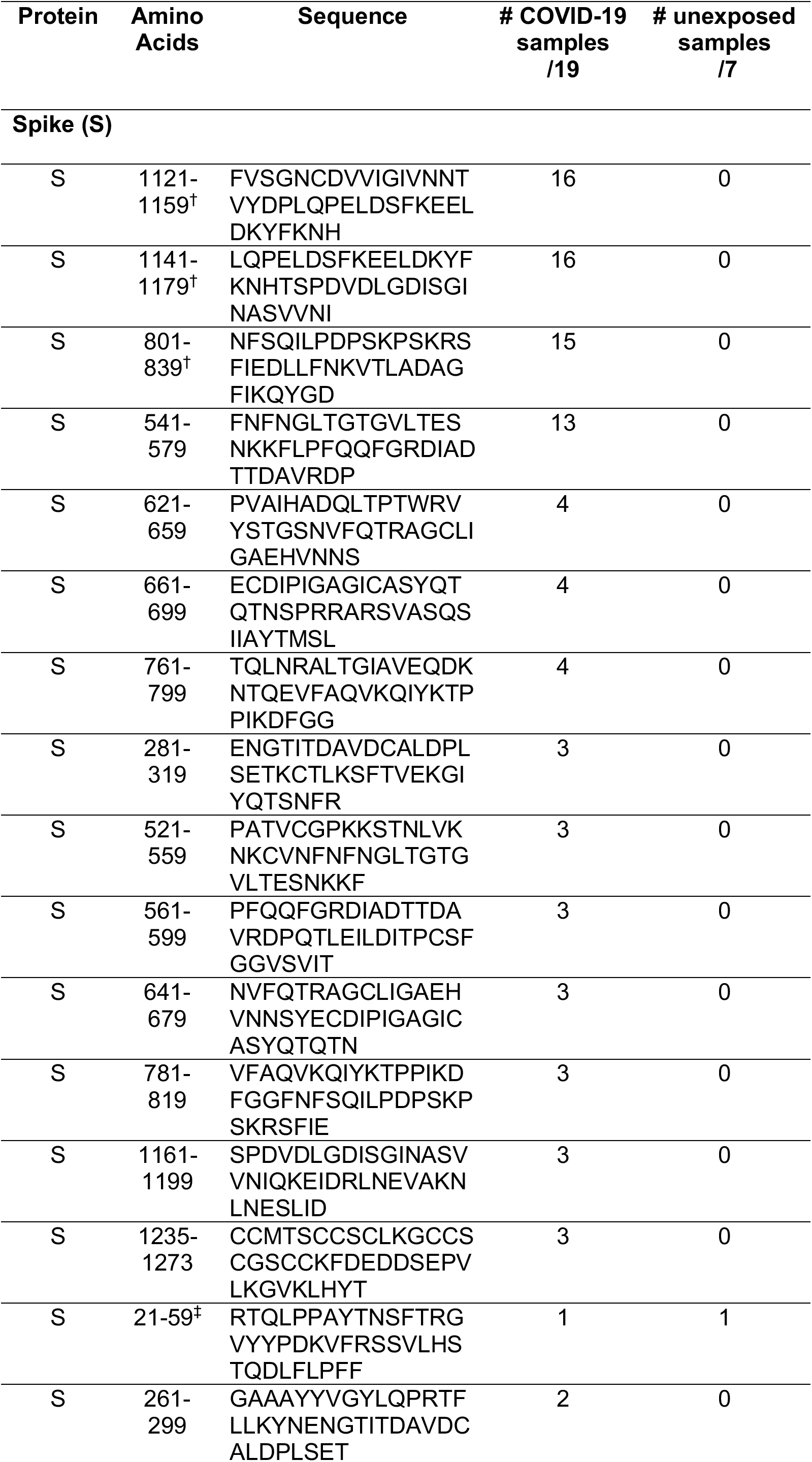
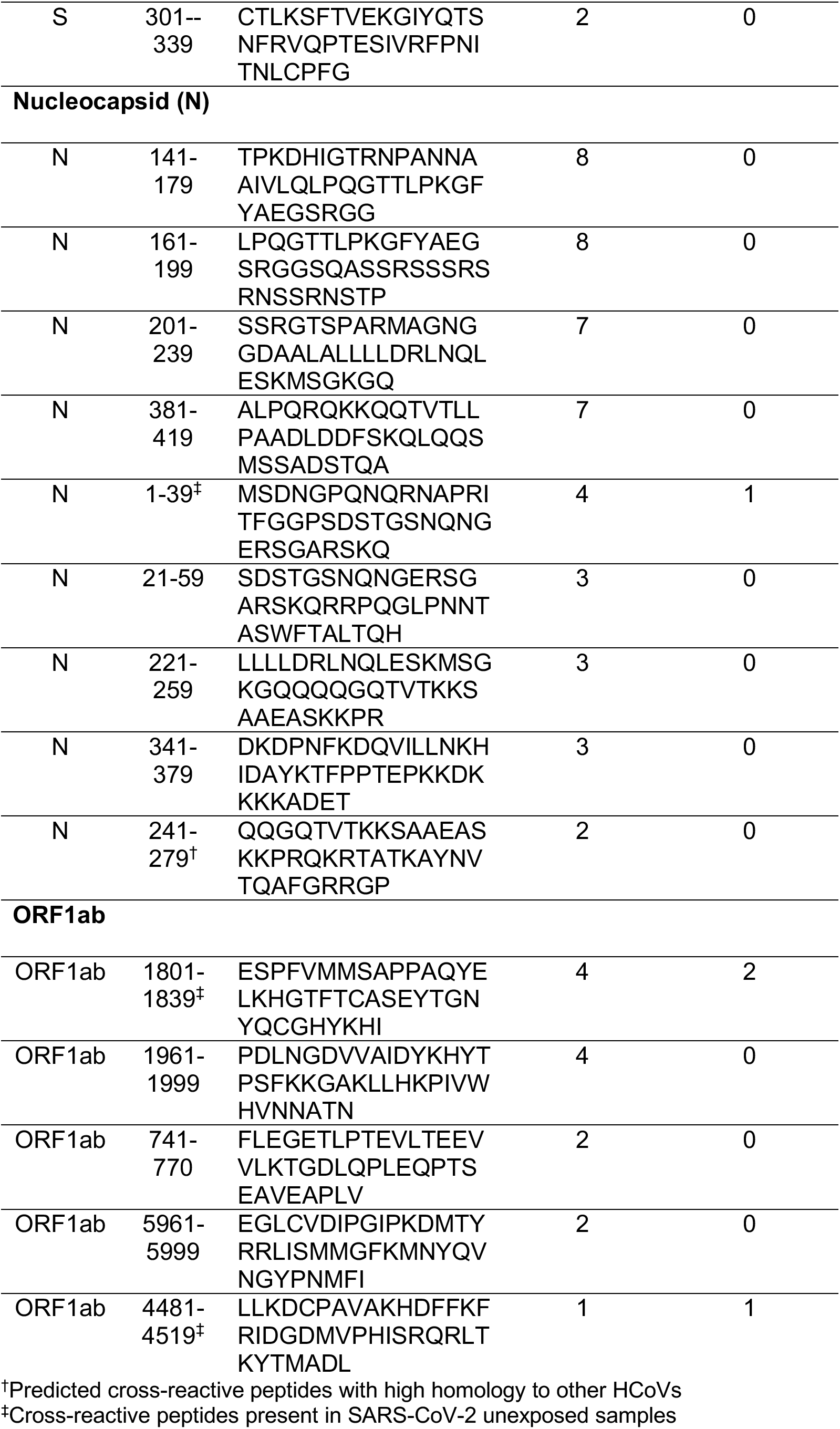
Top SARS-CoV-2 epitopes present in two or more individuals.

**Figure 4.**
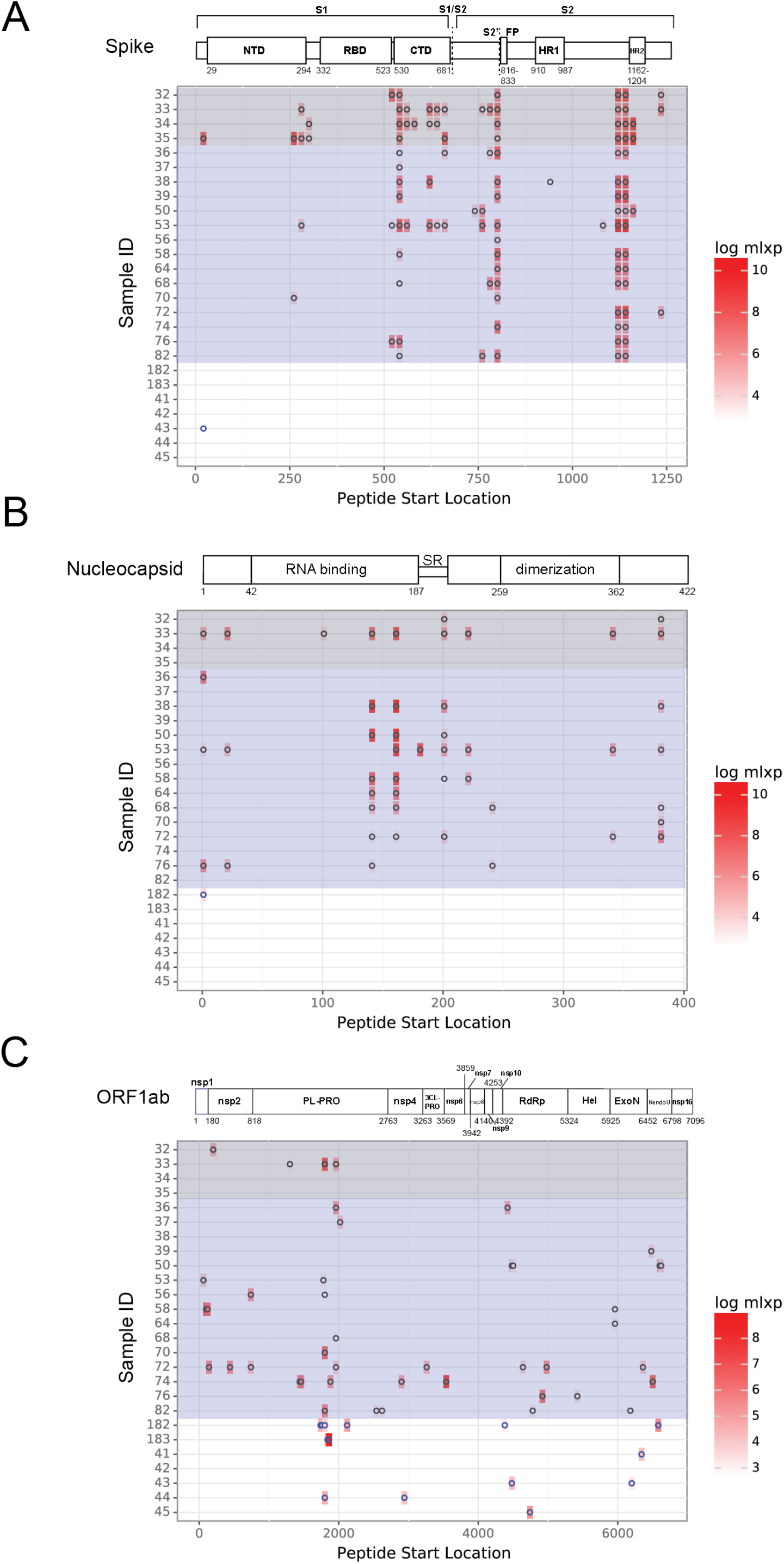
SARS-CoV-2 epitope profiles for immunodominant antigens. (**A-C**) Location of significantly enriched epitopes across the Spike protein (**A**), Nucleocapsid (**B**) and ORF1ab (**C**). Profiles for patients with COVID-19 are highlighted in grey (moderate/severe COVID-19) and purple (mild COVID-19). The remaining profiles are from SARS-CoV-2 unexposed individuals. Log(mlxp) values are indicated by the red gradient, shown to the right of the maps. Protein domain architecture for each antigen is above the heat map, with amino acid positions indicated.

Peptides derived from the N protein also elicited strong responses in many individuals. The most widely reactive region was N_141-199 (**Fig. 4B and Table 2)**. This region is composed of two overlapping peptides in the pan-CoV library, derived from the N protein RNA binding domain. Both peptides were reactive in eight individuals (six of which were identical for both overlapping peptides). We identified two additional sequences that were enriched across seven samples each: N_201-239, found upstream of the dimerization domain, and N_381-419, which is located downstream from the dimerization domain, at the C-terminus of the N protein. Five more epitopes that were reactive in four or fewer individuals are listed in **Table 2**.

The SARS-CoV-2 replicase polyprotein 1ab (ORF1ab) is composed of two overlapping reading frames that code for sixteen non-structural proteins, including the viral RNA-dependent RNA polymerase (RdRp), that are expected to be co- and post-translationally processed based on studies of SARS-CoV (Graham et al., 2008). We detected sequences with high mlxp values throughout ORF1ab, many of which were present in only a small fraction of individuals. The two most widely-reactive regions, ORF1ab_1801-1839 and ORF1ab_1961-1999, were both located within the papain-like protease (PL-PRO) sequence. Both of these sequences were reactive in four individuals with COVID-19. The remaining peptides identified from ORF1ab that were reactive in two or more individuals, including sequences from nsp2, exonuclease (ExoN) and the RdRp, are listed in **Table 2**. Lastly, we identified two peptides from the M protein (M_161-199 and M_181-219, which overlap by 19 aa), each present in one individual, and one peptide from ORF3a (ORF3a_237-274), present in one individual.

### Cross-reactive epitopes identified in pre-COVID-19 individuals and predicted based on cross-CoV sequence homology

Cross-reactivity in the viral antibody response can drive host immunity, complicate diagnostics and surveillance, and potentiate negative outcomes such as antibody-dependent enhancement. We sought to identify cross-reactive peptides between SARS-CoV-2 and other HCoVs by examining whether SARS-CoV-2 sequences were enriched in pre-pandemic, SARS-CoV-2 unexposed individuals. We identified four SARS-CoV-2 peptides that were reactive in at least one unexposed sample and at least one COVID-19 sample, suggesting that these were cross-reactive sequences likely arising from a prior HCoV infection (**Table 3**). One cross-reactive sequence found in the N protein (N_1-39) was detected in a sample with RT-PCR-confirmed prior endemic CoV infection, although the precise CoV species is unknown. We identified two cross-reactive peptides from ORF1ab (ORF1ab_1801-1839, from the PL-PRO protein and ORF1ab_6481-6520, from the RdRp protein), which shared ~35-47% and ~46-64% amino acid sequence identity with commonly circulating HCoVs, respectively (**Table 3**). Finally, we found a fourth cross-reactive peptide from the S protein, S_21-59, which shared only ~17% identity with HCoV-NL63, but 35.7% identity with HCoV-OC43 (**Table 3**). These peptides all exhibited significantly more homology with SARS-CoV and bat-SL-CoV, but these viruses are unlikely to have been the source of the antibody response because of the demographics associated with these individuals.

**Table 3.** Cross-reactive SARS-CoV-2 sequences from unexposed samples and their amino acid sequence homology with other CoVs.

| Protein | Amino acids | Total # patients<br>(# SARS-2 neg.) | % Identical (local alignment)* |  |  |  |  |  |  |
| --- | --- | --- | --- | --- | --- | --- | --- | --- | --- |
|  |  |  | OC43 | NL63 | HKU1 | 229E | MERS | SARS | Bat-SL |
| ORF1ab | 1801-1839 | 6(2) | 35.1 | 46.7 | 35.3 | 46.7 | 36 | 79.5 | 100 |
| N | 1-39 | 5(1) | 30.0 | 35.2 | 29.4 | 29.0 | 35 | 80.0 | 79.5 |
| S | 21-59 | 2(1) | 35.7 | 17.4 | 26.5 | 29.4 | 33.3 | 63.9 | 59.0 |
| ORF1ab | 4481-4519 | 2(1) | 63.2 | 46.2 | 64.1 | 59.0 | 66.7 | 92.3 | 89.7 |

\*Alignments to one representative strain of each HCoV

During our global fit, we identified 306 unique peptides arising from HCoVs other than SARS-CoV-2 in patients with COVID-19. These sequences were reactive because of a prior HCoV exposure, or because of cross-reactivity to antibodies generated by SARS-CoV-2 infection. To predict whether these sequences were positive in our analysis because of cross-reactivity with SARS-CoV-2 (measured by high levels of sequence conservation), we conducted local pairwise alignments across S, N and ORF1ab peptides from all HCoVs that were significantly reactive in at least two COVID-19 samples (**Fig. 5A)**. We used a Smith-Waterman alignment score of 55 as a cutoff for further investigation. We identified multiple peptides with high sequence similarity to SARS-CoV, as expected, given the higher genome-wide sequence similarity between SARS-CoV and SARS-CoV-2 (**Figs. 5B and S2**). We also isolated a subset of SARS-CoV-2 sequences with high homology to S and N sequences from two of the four commonly circulating endemic HCoVs, OC43 and HKU1. Local alignments between these sequences and the corresponding regions of SARS-CoV-2 revealed six- and seven-amino acid stretches of identity (**Fig. 5C**). Interestingly, none of the ORF1ab peptides that were significantly enriched in our study were highly conserved between SARS-CoV-2 and the other commonly circulating CoVs, despite the higher degree of conservation between HCoV ORF1ab sequences (**Fig. 5B**).

**Figure 5.**
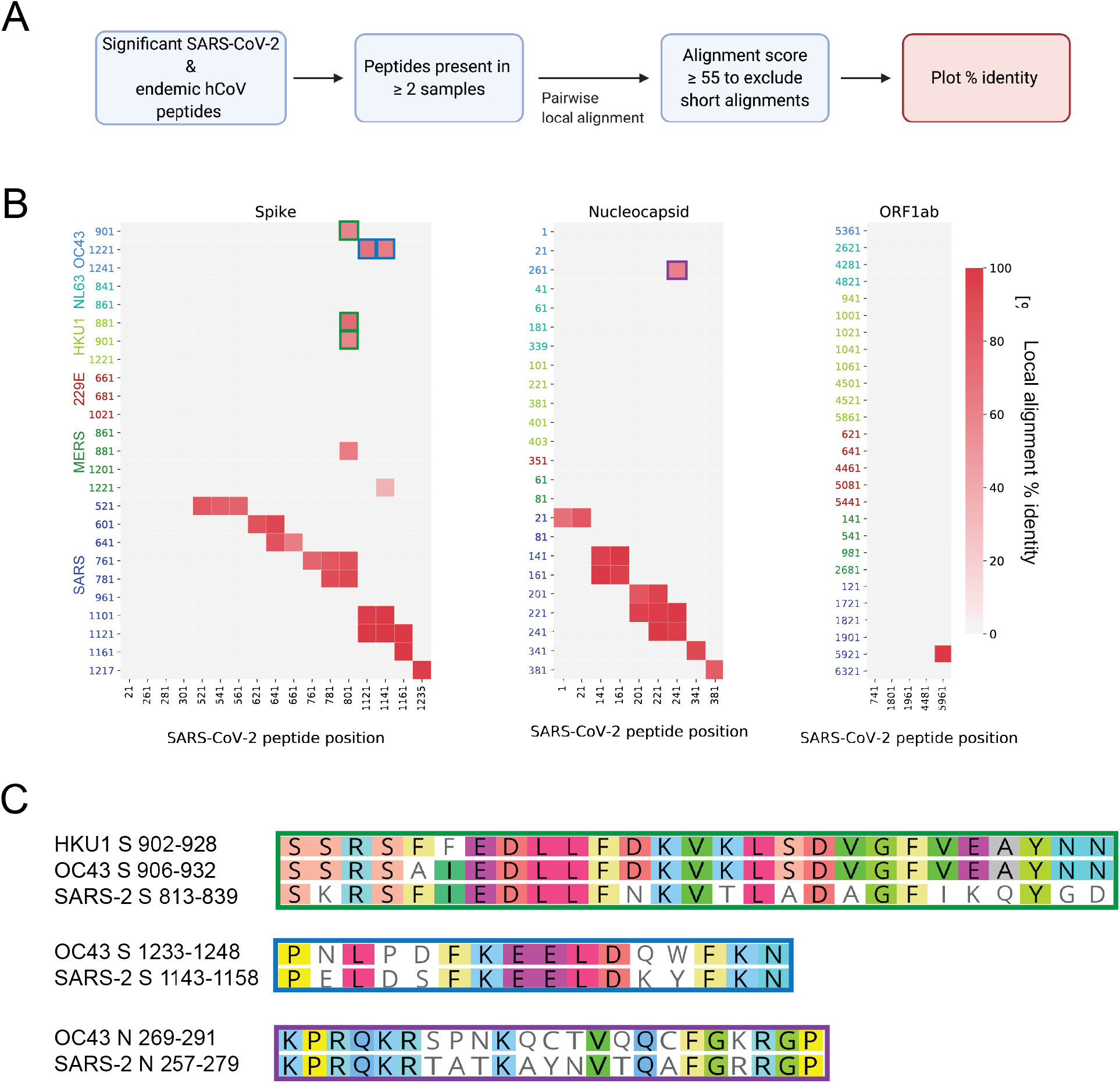
Identification of cross-reactive sequences among significant pan-CoV peptides. (**A**) Unique peptide hits from all CoVs that were present in two or more COVID-19 patient samples were aligned the Smith-Waterman method to probe for conservation. Sequences that were 100% identical between SARS-CoV-2 and the other CoVs were not included in the analysis. (**B**) Peptide pairs with alignment scores > 55 (Fig. S2) were plotted to show % identity. Peptide start positions from SARS-CoV-2 are listed on the x-axis and peptide start positions from the other human-infecting CoVs are listed on the y-axis. Green, blue and purple outlines match with the corresponding peptides pairs shown in (C). (**C**) Local sequence alignments for the high-scoring peptides in (B) from the commonly circulating HCoVs.

## Discussion

In this study we profiled the immune response to SARS-CoV-2 proteins in individuals with SARS-CoV-2 infections using phage display to capture linear immunogenic peptides spanning the entire proteome. By screening epitopes based on binding to SARS-CoV-2 protein sequences, we isolated epitopes with potential for both neutralizing and/or non-neutralizing activity. We identified S, N, and ORF1ab from SARS-CoV-2 as highly immunogenic and further isolated important regions at the epitope level. The proteome-wide capacity of our phage library and 39 amino acid tiling with 19 amino acid overlap resulted in a high-resolution and comprehensive definition of the targets of antibodies to SARS-CoV-2. SARS-CoV-2 epitopes stemming from the S protein were present in the highest density of patients with COVID-19. We identified 17 epitopes within the S protein that were present in two or more individuals, spanning both the S1 and S2 subunits, with some detected in > 75% of individuals. The breadth of antibody responses along the length of the S protein (and the other dominant ORFs) can inform interesting hypotheses about the SARS-CoV-2 immune response. For example, four individuals harbored antibodies targeting the S1/S2 cellular furin cleavage site, suggesting that this region of the S protein may be targeted when the SARS-CoV-2 virion is not yet mature (Hoffmann et al., 2020). Despite evidence for potently neutralizing antibodies targeting the S protein RBD, we did not identify epitopes in this region, possibly due to the tendency for RBD-directed antibodies to be conformational (Ju et al., 2020); such epitopes would only be detected with the phage display method if a large enough portion of the epitope was linear and not glycosylated. Finally, we found that epitopes from the S protein were dominant in moderate/severe COVID-19 samples versus mild COVID-19 samples, suggesting a correlation between COVID-19 severity and S protein epitope profile. This result coincides with recent evidence for stronger and more broad responses to both the S and N proteins in hospitalized COVID-19 patient samples (Shrock et al., 2020).

We identified nine epitopes within the N protein that were reactive in at least two individuals, four of which were present in at least 35% of patients. The two most reactive N protein epitopes were derived from the RNA binding domain. Epitopes derived from the non-structural N protein may be the results of exposure of the immune system to these antigens upon cell lysis and if they have activity, they are more likely to be non-neutralizing, owing to their sequestration away from the viral surface. However, their reactivity in our assay implies that the epitopes are accessible to antibody binding and may be useful in informing the design of new diagnostics. Epitopes isolated from ORF1ab, another non-structural polypeptide, were the most variable across patients. Of the 46 unique ORF1ab epitopes we identified, only five were present in two or more individuals, suggesting that ORF1ab responses are individual-specific. The two most reactive ORF1ab epitopes were situated in the PL-PRO sub-protein, an enzyme implicated in attenuating the host innate immune response and viral replication (Shin et al., 2020), and may be a target for therapeutic strategies that prevent viral replication.

Our pan-CoV phage library contained sequences from all seven human-infecting CoVs, allowing us to probe responses across the CoV family in COVID-19 and SARS-CoV-2 unexposed individuals. Our relatively small cohort of SARS-CoV-2 unexposed individuals was reactive to four “de facto” cross-reactive sequences from SARS-CoV-2. We also implemented a local alignment scheme to assess sequence homology to SARS-CoV-2 with reactive peptides from the other HCoVs and found three SARS-CoV-2 sequences with strong conservation to the commonly circulating HCoVs OC43 and HKU1. Additional studies using a larger cohort of SARS-CoV-2 unexposed individuals or individuals with specific endemic CoV diagnoses will be necessary to further evaluate these epitopes. These highly homologous sequences, along with the cross-reactive sequences from unexposed individuals, may be valuable in future diagnostic and surveillance efforts aimed at discrimination of HCoV serological profiles, and may inform studies assessing the risk of antibody-dependent enhancement.

Our study included a relatively small sample size, which did not allow us to draw robust conclusions regarding the frequency of responses or whether there were differences in the response based on disease severity. Nevertheless, in light of the ongoing SARS-CoV-2 pandemic, our data reveal viral proteome-wide antibody binding signatures in patients with confirmed COVID-19. Given the urgency for targeted SARS-CoV-2 vaccine and therapeutic development, and the importance of prospective surveillance to detect the emergence of potential antibody escape variants, it is absolutely essential that epitope mapping studies be validated using multiple approaches. Indeed, our results nicely converge with a number of other studies aimed at mapping the epitope profiles of SARS-CoV-2 and together begin to provide a comprehensive picture of the responses to this pandemic virus (Amrun et al., 2020; Poh et al., 2020; Shrock et al., 2020). A recent study using a similar phage immunoprecipitation approach with a larger virus library and different analytical techniques reassuringly identified the same three main protein targets of the SARS-CoV-2 antibody response, and many epitopes that overlap with those identified in this study, despite the divergence in methodology (Shrock et al., 2020). Our phage-based profiling method, coupled with robust computational modeling of significantly enriched sequences provides an important launch point for further characterization of neutralizing, non-neutralizing, and cross-reactive antibodies targeting SARS-CoV-2.

## Materials and Methods

### Samples

In this study, all patients with mild COVID-19 were outpatients, not requiring hospitalization. Patients with moderate/severe COVID-19 were hospitalized and a range of clinical outcomes were documented, ranging from supplemental oxygen, intubation and death. All samples were heat-inactivated at 56 °C for 60 minutes prior to short-term storage at 4°C or long-term storage at −80°C. An additional two endemic CoV positive, and three mild COVID-19 samples were tested in the phage display assay but were not included in the global fit because of poor in-assay technical replicate correlation or poor correlation between experiments conducted using Library 1 and Library 2.

### Pan-CoV phage display library construction and immunoprecipitation

Epitope mapping via phage display and immunoprecipitation was carried out essentially as previously described (Mohan et al., 2018; Williams et al., 2019). An oligonucleotide pool was generated from 17 CoV protein coding sequences retrieved from GenBank: OC43-SC0776 (MN310478), OC43-12689/2012 (KF923902), OC43-98204/1998 (KF530069), 229E-SC0865 (MN306046), 229E-0349 (JX503060), 229E-932-72/1993 (KF514432), NL63-ChinaGD01 (MK334046), NL63-Kilifi_HH-5709_19-May-2010 (MG428699), NL63-012-31/2001 (KF530105), NL63-911-56/1991 (KF530107), HKU1-SI17244 (MH90245), HKU1-N13 genotype A (DQ415909), HKU1-Caen1 (HM034837), MERS-KFMC-4 (KT121575), SARS-Urbani (AY278741), SARS-CoV-2-Wuhan-Hu-1 (MN908947), bat-SL-CoVZC45 (MG772933). Multiple strains of the endemic CoVs were selected to cover a wide range of circulation chronology using the Nextstrain database (Hadfield et al., 2018). A single HIV-1 envelope sequence was also included for controls (BG505.W6.C2, DQ208458).

Bacterial codon-optimized oligonucleotide libraries were designed using the Python script available at https://github.com/jbloomlab/phipseq_oligodesign. During the design process, viral protein coding sequences were reverse translated to DNA in 39 amino acid tiles with 19 amino acid overlaps. Adaptor sequences (5’: AGGAATTCTACGCTGAGT and 3’: TGATAGCAAGCTTGCC) were added. Two separate oligonucleotide pools with equivalent design were commercially synthesized (Twist Biosciences). The libraries were PCR amplified using in-house primers (FWD: AATGATACGGCAGGAATTCTACGCTGAGT and REV: CGATCAGCAGAGGCAAGCTTGCTATCA), digested, cloned into the T7Select 10-3b Vector, packaged in T7 phage and amplified according to manufacturer instructions (EMD Millipore).

For phage immunoprecipitation, 1.1 mL 96-deep-well plates (CoStar) were blocked with 3% BSA in TBST (Tris-buffered saline-Tween) by rocking overnight at 4°C. Amplified phage library was diluted in Phage Extraction Buffer (20 mM Tris-HCl, pH 8.0, 100 mM NaCl, 6 mM MgSO_4_) to reach 2×10^5^-fold phage representation (1.33×10^9^ PFU/mL for a phage library containing 6,659 sequences) and added to each well at a volume of 1 mL. We estimated plasma and serum IgG concentrations to be 10 ug/uL (Mabuka et al., 2012) and added 10 ug of each sample to the diluted phage library in duplicate for a total of two technical (within-assay) replicates per experiment. Serum/plasma antibodies were allowed to bind to the phage library by rocking at 4°C for 20 hours. To account for non-specific interactions during the immunoprecipitation step, we prepared multiple wells with no serum and only phage library (“mock”-immunoprecipitations) and treated them to the same rocking procedure. To immunoprecipitate phage-antibody complexes, 40 uL of a 1:1 mixture of protein A and protein G magnetic Dynabeads (Invitrogen) was added to each well and rocked at 4°C for 4 hours. Dynabeads were then isolated using a magnetic plate, washed three times in 400 uL of wash buffer (50 mM Tris-HCl, pH 7.5, 150 mM NaCl), 0.1% NP-40), and resuspended in 40 uL of water. Dynabead-bound phage were lysed by incubating samples at 95°C for 10 minutes and stored at −20°C prior to Illumina library preparation.

### Illumina library preparation

Phage DNA was PCR-amplified with Q5 High-Fidelity DNA polymerase (New England Biolabs) in two rounds to produce Illumina libraries containing adaptor sequences and barcodes for multiplexing. Round 1 PCRs were performed using 10 uL of resuspended, lysed phage in a 25 uL reaction volume using primers R1_FWD (TCGTCGGCAGCGTCTCCAGTCAGGTGTGATGCTC) and R1_REV (GTGGGCTCGGAGATGTGTATAAGAGACAGCAAGACCCGTTTAGAGGCCC). Round 2 PCRs were performed using 2 uL of the Round 1 reaction in a 50 uL final volume with unique dual-indexed primers as previously described (Williams et al., 2019). Round 2 PCR products were quantified using Quant-iT PicoGreen according to manufacturer instructions (Thermo Fisher). Samples were pooled in equimolar quantities, gel purified, and submitted for sequencing on a MiSeq with 126 bp single-end reads.

### Phage sequence alignment pipeline

An enrichment matrix was created by aligning all sequences to the pan-CoV reference library using a Nextflow data processing pipeline (https://github.com/matsengrp/pan-CoV-manuscript) (Di Tommaso et al., 2017). The pipeline was initiated with metadata for all samples (including a path to the fastq reads) as well as the metadata for all peptides in the library. The processing steps were as follows: (1) We built a Bowtie index from the peptide metadata by converting the metadata to fasta format and feeding it into the bowtie-build command (Langmead et al., 2009). (2) We aligned each of the samples to the library using end-to-end alignment allowing for up to two mismatches. Each read was 125 base pairs long, and the low-quality end of the read was trimmed to match the reference length, 117 base pairs, before alignment. (3) We extracted the peptide counts for each sample alignment using samtools-idxstats (Li et al., 2009). (4) All individual counts information for each sample were merged into an enrichment matrix. The resulting dataset containing the enrichment matrix, sample metadata, and peptide metadata was organized using the xarray package (Hamman and Hoyer, 2017).

### Assessment of peptide significance using a Gamma-Poisson model

To determine significance of enriched peptides in the background of noise introduced by non-specific immunoprecipitation, curated sample sets from two separate phage libraries were fit to a Gamma-Poisson mixture model in the phip-stat Python package provided by the Laserson Lab (https://github.com/lasersonlab/phip-stat). For simplicity, we focused our downstream analyses on one strain from each of the CoV species included in the phage libraries. We required samples to have (1) high technical (in-assay) correlation and (2) high correlation in experiments conducted with phage libraries 1 and 2, in order to be included in the fit. For each model fit, data from mock-IP controls were included with the patient sample data to better account for the abundance and non-specific binding associated with each peptide in the phage library. To control for the variance in sequencing coverage between samples, we first normalized all samples using counts factor method (Anders and Huber, 2010). This resulted in a normalized raw counts matrix, M, with *i* peptides and *j* samples. The model assumes that each entry in the count matrix, for any given peptide *i*, is sampled from a Poisson distribution with rate, λ_i_. Next, we assumed the prior distribution of any λ_i_ is a Gamma distribution defined by α and β parameters. We used the scipy.optimize package (Virtanen et al., 2020) to infer maximum likelihood estimates that would generate a set of mean normalized counts values across samples for each peptide, *i*. Given that the posterior of the rate is also Gamma distributed, the posterior hyperparameters for each peptide, *i*, are given by the formulas 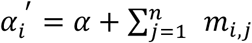 and 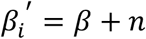, where *n* is the number of samples. Because the gamma distribution is a conjugate prior for the Poisson, we get 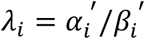 for each peptide. Finally, −log_10_(pval) (mlxp) values are by computed using the value of the tail of the Poisson distribution for each normalized sample count at peptide, *i*.

### False positive rate (FPR) estimation using HIV Env peptides

We observed quite extreme p-values using the Gamma-Poisson approach (as well as using the Generalized Poisson approach in (Larman et al., 2011)), indicating that these p-values were not well calibrated. Thus, the selection of peptides with significant binding affinity was performed by applying a minimum threshold requirement on the mlxp, which was set based on using HIV peptides as a control. Specifically, peptides derived from the HIV-1 envelope were presumed not to truly bind with SARS-CoV-2 antibodies, so we used these HIV-1 peptides to estimate the FPR for a threshold under consideration: the number of HIV-1 sample-peptide pairs above this threshold divided by the total number of HIV-1 sample-peptide pairs in the library batch after the curation step. Hence, for each library batch, we set the threshold to be the value where 5% of the HIV-1 peptides have mlxp values above the threshold. In each library batch, there were a total 798 HIV-1 sample-peptide pairs. None of the HIV peptides have significant sequence homology with SARS-CoV-2 peptides.

### Local sequence alignment

The similarity between SARS-CoV-2 and other CoV peptides was quantified by performing local alignment and then computing the identical fraction: the fraction of matching amino acids at each position of the aligned subsequence. The Smith-Waterman algorithm was applied with the BLOSUM62 cost matrix, a gap open penalty of 12, and a gap extension penalty of 3 (Smith and Waterman, 1981). We used the pairwise2 function of the Biopython software package to perform the alignment (Cock et al.).

## Acknowledgments

We extend gratitude to all study participants who contributed samples. We thank members of the Overbaugh, Matsen and Chu labs for thoughtful discussion and advice. We thank Trevor Bedford (FHCRC) for advice on strains to include in the pan-CoV phage library and Theodore Gobillot (FHCRC) for advice on phage library construction and immunoprecipitation. This work was funded by NIH grants AI138709 (PI Overbaugh), and R01 AI146028 and U19 AI128914 (PI Matsen). Julie Overbaugh received support as the Endowed Chair for Graduate Education (FHCRC). The research of Frederick Matsen was supported in part by a Faculty Scholar grant from the Howard Hughes Medical Institute and the Simons Foundation.

## Supplementary figures

**Table S1.** Additional demographic information.

|  | **Mild (n = 14)** | **Moderate/Severe (n = 5)** | **Endemic (n = 2)** |
| --- | --- | --- | --- |
| **Mean Age (Mean ± SD)** | 42.2 ± 14.6 | 62.9 ± 7.3 | 64 ± 3.0 |
| **Female (n, %)** | 8 (57.1) | 1 (20.0) | 0 (0.0) |
| **Race/ethnicity (n, %)** |  |  |  |
| White, non-Hispanic/Latino | 12 (85.7) | 2 (40.0) | 2 (100.0) |
| African American, non-Hispanic/Latino | 0 (0.0) | 0 (0.0) | 0 (0.0) |
| Other, non-Hispanic/Latino | 1 (7.1) | 3 (60.0) | 0 (0.0) |
| Hispanic/Latino | 1 (7.1) | 0 (0.0) | 0 (0.0) |
| **Insurance Status (n, %)** |  |  |  |
| Public (or none/self-pay) | 2 (14.3) | 3 (60.0) | 2 (100.0) |
| Private or both | 12 (85.7) | 2 (40.0) | 0 (0.0) |
| **Number of underlying conditions (n, %)** |  |  |  |
| $\geq 1$ | 0 (0.0) | 3 (60.0) | 2 (100.0) |
| None | 14 (100.0) | 2 (40.0) | 0 (0.0) |

**Figure S1.**
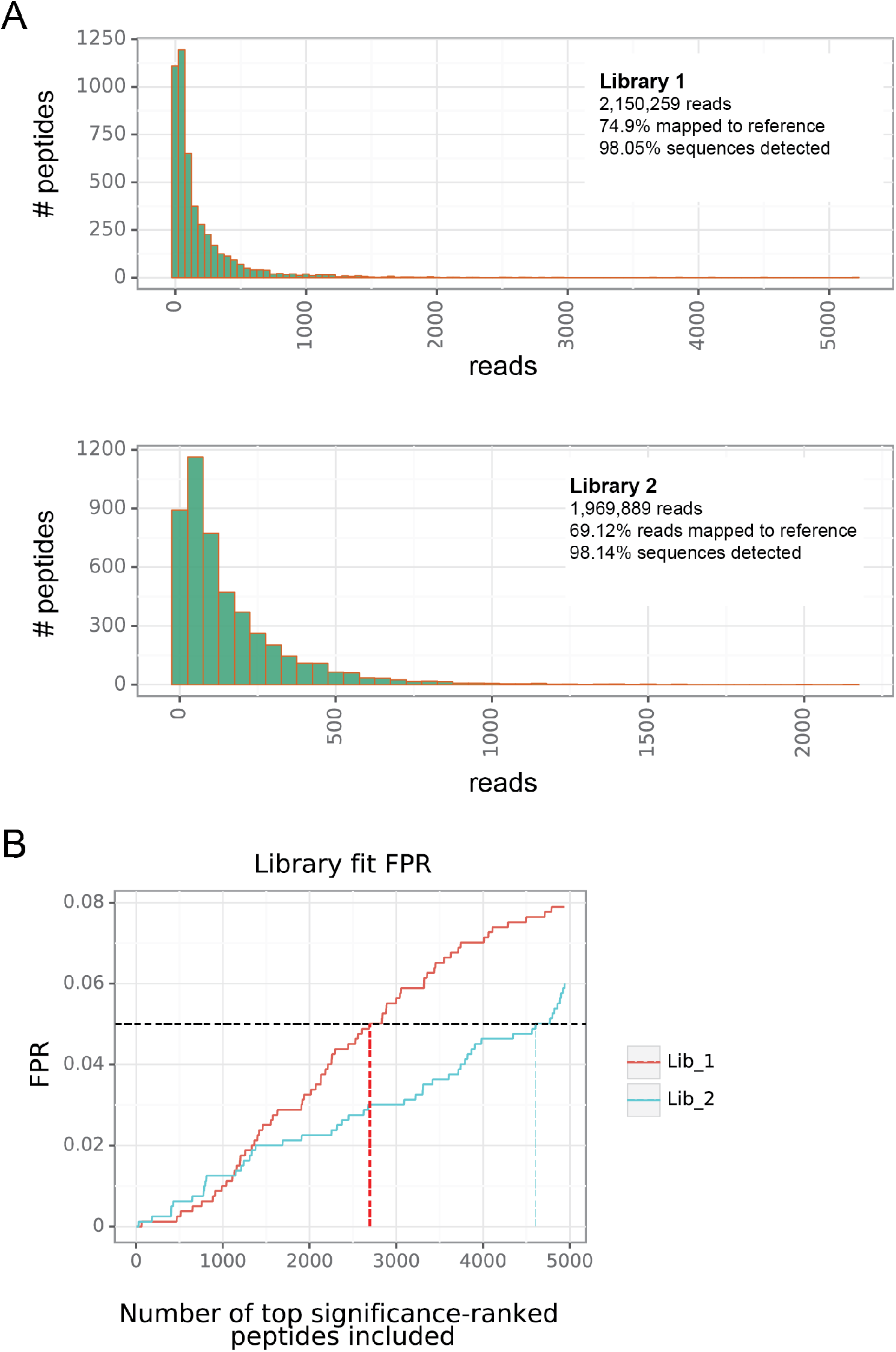
Pan-CoV input phage library coverage and FPR modeling for significance thresholding. (**A**) Number of peptides plotted against number of reads for both independently generated phage libraries. Text inlays show the total number of reads sequenced from the input libraries, the percentage of reads mapping to the target CoV peptide sequences, and the percentage of peptide sequences with zero reads. (**B**) The number of peptides included in the analysis plotted against the FPR, as calculated from the presence of enriched HIV-1 sequences. A FPR of 0.05 was chosen as a threshold and the number of peptides this cutoff corresponded to is indicated by red and blue lines for Libraries 1 and 2, respectively.

**Figure S2.**
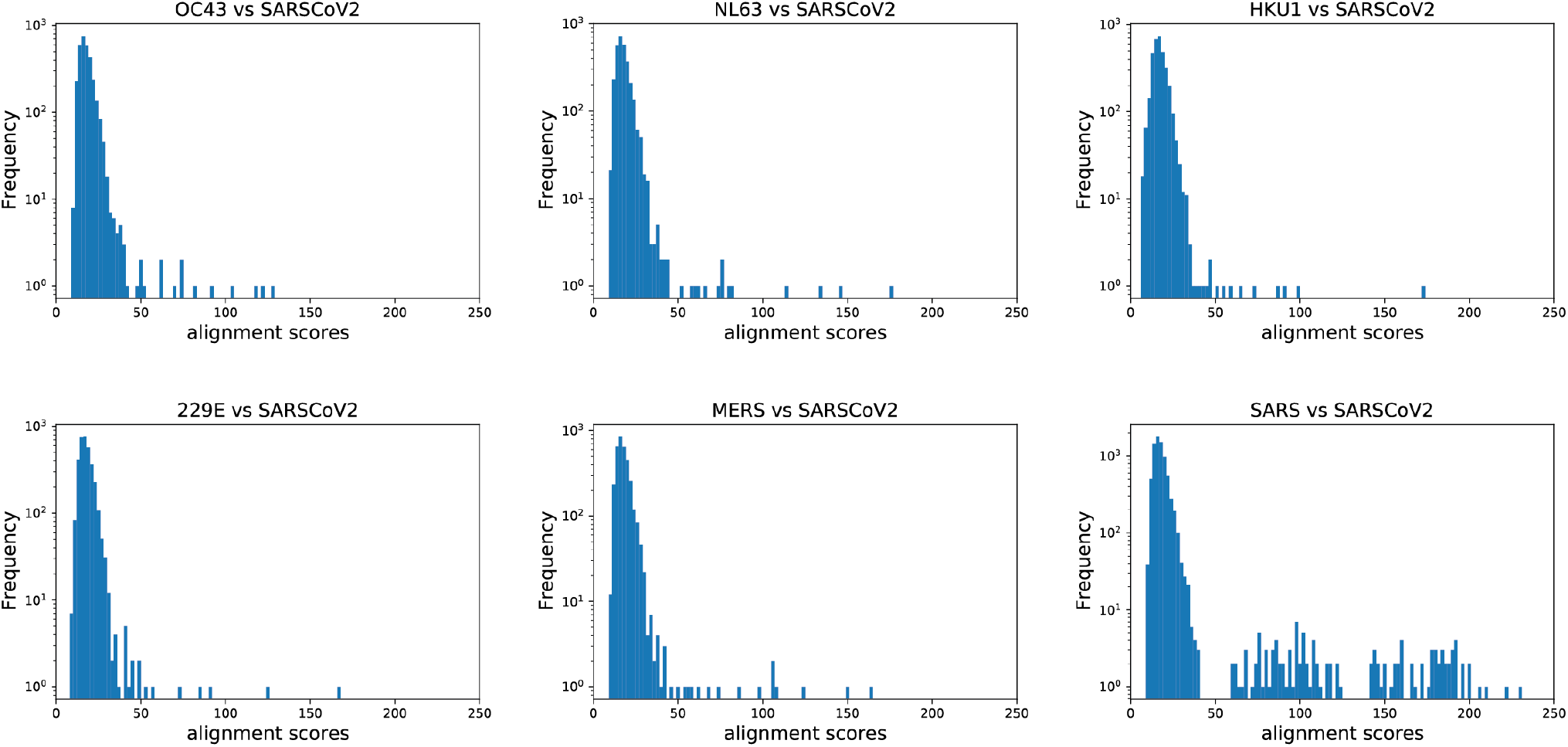
Smith-Waterman alignment scores for significant epitopes from all HCoVs. Pairwise local alignment scores between significant SARS-CoV-2 and other commonly circulating or pathogenic HCoV peptides plotted against peptide pair frequency.

